# Phylogeny and disease links of a widespread and ancient gut phage lineage

**DOI:** 10.1101/2023.08.29.555303

**Authors:** Patrick A. de Jonge, Bert-Jan H. van den Born, Aeilko H. Zwinderman, Max Nieuwdorp, Bas E. Dutilh, Hilde Herrema

## Abstract

Viruses are a core component of the human microbiome, impacting health and disease through interactions with gut bacteria^1^ and the immune system^2^. Most viruses in the human microbiome are bacteriophages, which exclusively infect bacteria. Individual gut bacteriophages can affect bacterial bile acid deconjugation^3^, and can alter their infection strategy based on dietary content^4^. Up to recently, most studies of the gut virome have focused on low taxonomic scales (e.g., viral operational taxonomic units), hampering population-level analyses. We previously identified the expansive and widespread Candidatus *Heliusviridae* bacteriophage family in a cohort with inhabitants of Amsterdam, the Netherlands. Here, we study their biodiversity and evolution in a wide variety of human populations. With a detailed phylogeny based on sequences from six viral genome databases, we now propose the Candidatus order *Heliusvirales* to accommodate these viruses. We identify *Ca. Heliusvirales* viruses in 80% of 5,441 individuals across 39 studies, and also in nine out of thirteen analyzed metagenomes from ancient humans that lived in Europe and North America between 1,000 and 5,000 years ago. We show that a large *Ca. Heliusvirales* lineage has diversified starting at the appearance of *Homo sapiens* some 200,000-300,000 years ago. Ancient peoples and modern hunter-gatherers further have distinct *Ca. Heliusvirales* populations that are characterized by lower richness than modern urbanized people. Within urbanized people, those suffering from type 1 and type 2 diabetes, as well as inflammatory bowel disease, have higher *Ca. Heliusvirales* richness than healthy controls. We thus conclude that these ancient core members of the human gut virome have thrived with increasingly westernized lifestyles of the human population.

## Introduction

Bacteriophages are the second most common biological entities in the human gut microbiome, after their bacterial targets^5^. As obligate parasites of bacteria, phages play a key role in regulating the structure and function of the gut microbiome in health and disease^6^. For example, phage community alterations have been linked to diseases such as inflammatory bowel disease^1,7^, metabolic syndrome^8^, and malnutrition^9^. Individual human gut phages have further been found to mediate diet-induced bacterial lysis^4^, bacterial bile acid metabolism^3^, and gut colonization^10^. But while phages are evidently crucial to the functioning of the gut microbiome, their study at higher taxonomic levels was long restricted to morphology-rather than phylogeny-based taxonomy.

This knowledge gap is partly due to the absence of a universal viral marker gene akin to ribosomal RNA in cellular organisms. Bacteriophage taxonomy was originally based on nucleic acid composition (with the majority being dsDNA viruses) and viral particle morphology^11^, with the bulk of known bacteriophage diversity being classified as *Siphoviridae, Myoviridae,* or *Podoviridae*. These families, as well as the order *Caudovirales*, proved to be so genomically diverse that they were recently abolished and reclassified in the class *Caudoviricetes*^12,13^. A striking example of the discovery of higher taxonomic lineage was the highly prevalent and abundant human gut phage crAssphage^14^ and its relatives^15^ in metagenomics data. These viruses are now accepted as the phylogenetics-based *Crassvirales* order by the International Committee for the Taxonomy of Viruses (ICTV). But while the description of the *Crassvirales* is a step forward in the taxonomy of the human gut virome, many other important lineages remain unclassified.

We recently identified the putative phage family *Ca. Heliusviridae* in a study of gut virome perturbations associated with metabolic syndrome, a collection of risk factors for cardiometabolic disease^8^. The expansive family was detected in over 90% of 196 study participants and may thus be part of a persistent core of human gut bacteriophage lineages^16,17^. But as our previous study focused on a single cohort, it remains unknown how widespread these phages are in the general population, as well as in other environments. Furthermore, their relation to human health beyond metabolic syndrome is yet to be elucidated. Their further adoption in virome studies necessitates a robust taxonomic classification based on comprehensive phylogenetic analysis of sequences from diverse sources^18–21^.

Here, we present a comprehensive study of *Ca. Heliusviridae* genomic and phylogenetic diversity. Comparisons of member phages derived from several large metagenomic databases show both the distinctness of this family from known ICTV taxonomic lineages, and their phylogenetic structure. We further provide evidence that increasing ecological richness of this phage family in the gut is linked to both urbanized lifestyles and several (cardiometabolic) diseases.

## Results

### Helius-phages are ubiquitous in bacteriophage databases

We previously reported an expansive and widespread clade of human gut phages, tentatively named as the family *Ca. Heliusviridae*, after the HELIUS (Healthy Life In an Urban Setting) cohort that we studied^8^. These phages shared nine marker genes: four structural proteins (terminase large subunit or TerL, portal protein, major capsid protein, and a head maturation protease), three replication-related ones (DNA polymerase I, helicase, and nuclease), and two proteins of unknown function (**Figure 1a**)^8^. To study these phages more extensively, we analyzed 842,163 contigs of over 30 kbp from seven phage databases with profile hidden Markov models (HMMs) of all nine marker genes (**Figure 1b**). Among the 249,366 contigs that were deemed complete by checkv^22^, 33,356 had a hit against the terminase large subunit (TerL) marker gene. The TerL, which encodes an ATP-driven molecular motor involved in genome translocation into the viral capsid, is highly conserved among *Caudoviricetes* phages and consequently frequently used as marker gene^13,15^. Contigs with a bit-score of over 700 against the TerL tended to have a hit against at least four other marker genes, and those with a hit against all marker genes always had a TerL bit-score of over 700 (**Figure 1c**). We hypothesized that a bit-score cutoff of 700 against the TerL separates *Ca. Heliusviridae* from other phages. To test this, we built a tree of TerL genes from all 1,290 contigs that passed this threshold and representative genomes from International Committee on the Taxonomy of Viruses (ICTV)-recognized virus lineages with a TerL bit-score of ≥50. The highest TerL bit-score of an ICTV-recognized virus was 515.4 against *Dragolirvirus dragolir* (*Gochnauervirinae,* MG727697). The resulting maximum-likelihood tree clearly showed a separation between ICTV-recognized viral lineages and the prospective *Ca. Heliusviridae* (**Figure 1d**). The exception were five sequences that had TerL bit-scores of between 700 and 760, and further only carried the portal protein and major capsid protein. This might have resulted from transferal of capsid proteins between phages. For further study, we selected the 1,285 complete viral genomes that comprised the distinct branch in the tree. Phage-specific gene annotation with pharokka^24^ refined the function of several marker genes: the DNA polymerase I was revealed to instead be an exonuclease, while the ones of unknown function encode a DNA polymerase and ssDNA-binding protein (**Figure 1a**). We next determined which genes were most common among these phages by forming clusters of homologous proteins (i.e., protein clusters). The only universal protein cluster was the TerL, and the next eight most common protein clusters represented the other marker genes (**Figure 1e**, **Supplementary Table 2**). Genes with hits against the HMMs of the major capsid protein, head maturation protease, helicase, and DNA polymerase genes were spread out over multiple protein clusters, indicating some heterogeneity in their sequences. Overall, all nine marker genes were found in at least 1000 of the genomes (**Supplementary Table 3**). Thus, the nine originally identified marker genes are extremely common, though not universal, among *Ca. Heliusviridae*.

**Figure 1:**
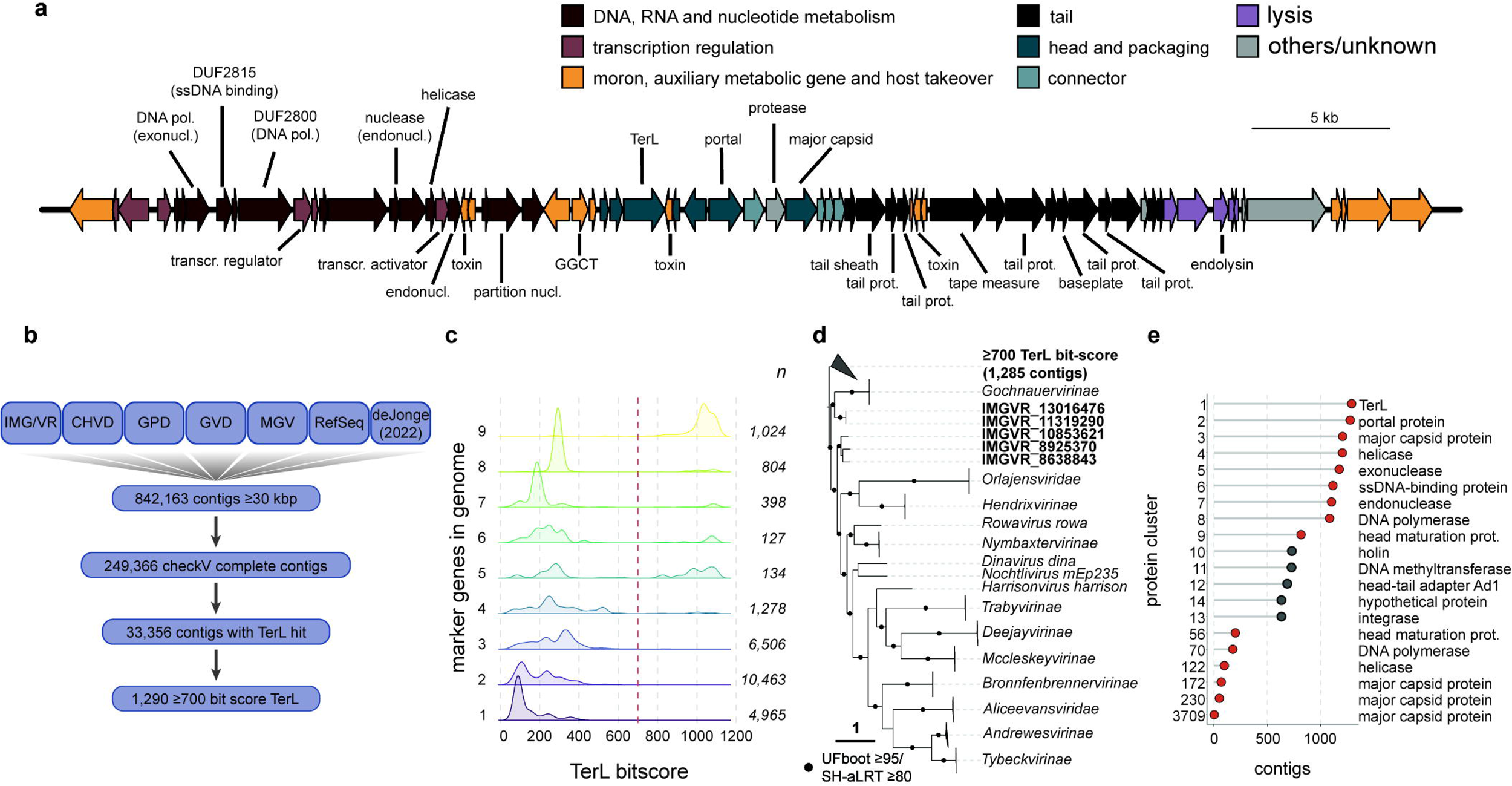
Presence of near-universal marker genes is linked to TerL homology. **a:** Annotation of NatCom_15117, a complete genome with inverse terminal repeats. The nine marker genes are denoted above the genome line, with re-annotated functions in parentheses. Other annotated genes are below the genome line. Full annotation is reported in **Supplementary table 1. b:** Overview of the procedure by which genomes were selected from seven phage databases. **c:** density plots of bit-scores from a search with the TerL HMM, separated by the total amount of marker genes found in the genome. Dots are individual complete genomes. **c:** a rooted approximate maximum-likelihood tree of TerL genes with hmm bit-scores over 700 and members of all ICTV lineages with similar TerL genes (hmm bit-score ≥50). Groups are collapsed, for un-collapsed version of the tree, see Figure 2a. **e:** prevalence of the most common protein clusters. Red denotes marker genes, grey others.

### Sub-taxa with distinct genomic characteristics reflect ecologically divergent Helius-virus lineages

We next sought to establish more detailed phylogenies of these phages. We previously divided the *Ca. Heliusviridae* into three groups: *alpha, beta,* and *gamma*. In an approximate maximum-likelihood tree of TerL genes, we again discerned three large well-supported clades (Shimodaira–Hasegawa-like approximate likelihood ratio test^25^ ≥80 and ultrafast bootstrap approximation^26^ ≥95, **Figure 2a**), but these did not completely equate to the previous putative groups. The largest clade contained all but one of the partial genomes from the previously proposed *gamma*-group, but groups *alpha* and *beta* were spread out across the other two clades. This reflects the two main clades in group *beta* that were evident in the earlier analysis^8^, while their different architecture in the TerL-based tree versus the earlier concatenated tree likely reflects horizontal gene transfer of at least some of the core genes among these groups. Furthermore, large subclades in all three clades contained no representative of the earlier groups, and neither did a number of smaller clades across the tree which were not part of the three main clades. In the face of this expanded biodiversity, we set out to draw more detailed taxonomic boundaries to combine related Helius-virus lineages, allowing for ecological and evolutionary interpretations.

**Figure 2:**
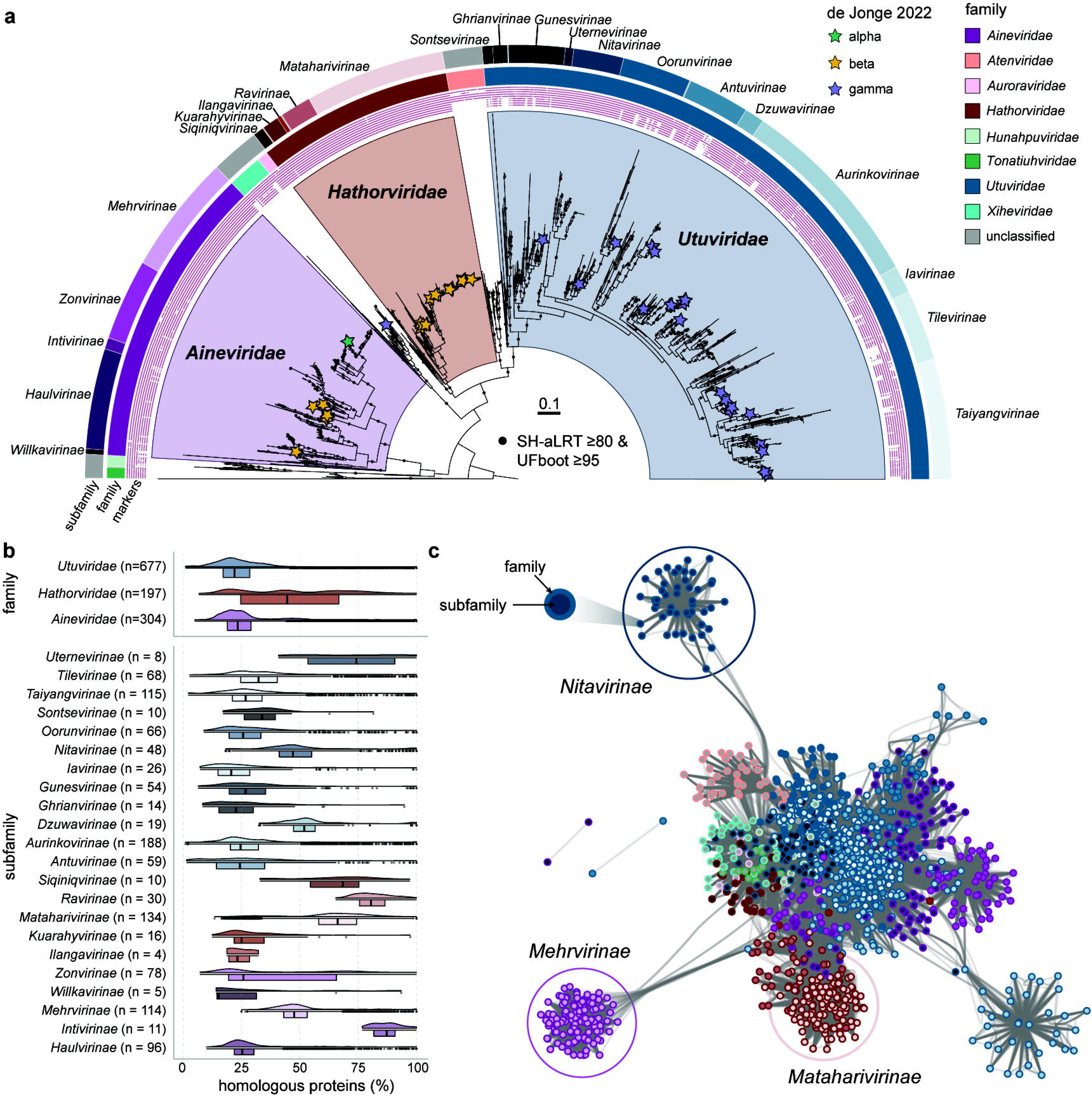
The *Ca. Heliusvirales* are a large order of gut phages with distinct subgroups. **a:** a rooted approximate maximum-likelihood tree of *Ca. Heliusvirales* TerL genes. The outgroup at the extreme left of the tree are the ICTV-sequences depicted in Figure 1d. The symbols for the presence of the marker genes are, from inside to outside: the exonuclease, ssDNA binding protein, DNA polymerase, endonuclease, helicase, TerL, portal protein, head maturation protease, and major capsid protein. **b:** chart depicting the pairwise percent of shared genes among the three main families (top) and their subfamilies (bottom). The n reflects the number of genomes in the lineage. For an overview of all families, see **Supplementary Figure 1**. Box plots show the median (middle line), 25th, and 75th percentile (box), with the 25th percentile minus and the 75th percentile plus 1.5 times the interquartile range (whiskers), and outliers (single points). **c:** vContact2 protein-sharing network of all sequences in **a**. Node edge color denotes family, while fill denote subfamilies, both according to the same colors as use in **a**.

The TerL tree also made it evident that these phages are better classified at the order-level. The reasoning behind this is as follows: the three largest clades were divisible into smaller clades that at minimum shared 5-57% of their protein content, too low for them to be considered different viral genera (**Figure 2b, Supplementary Figure 1**)^27^. In line with the viral taxonomic hierarchy^28^, we classify these clades as subfamilies. The higher-level clades are then necessarily families, and the entirety of the tree an order or suborder. Since suborders are strictly optional levels, and there is no clear distinction between them and orders^28^, we tentatively reclassify the phages studied here at the order level. Consequently, we renamed the Ca. *Heliusviridae* as the Candidatus viral order *Heliusvirales.* Because Helios was the ancient Greek solar deity, we named families after various solar deities. The three largest families are the *Ca. Utuviridae* (Mesopotamian), *Ca. Aineviridae* (Irish), and *Ca. Hathorviridae* (Egyptian). Several smaller well-supported and high-level clades were also placed at the family rank due to their very low (<15%) shared protein content (**Supplementary Figure 1)**. The subfamilies within the three largest families, mostly had a median shared protein content of 25% (**Figure 2b)**. All subfamilies were named after words for sun in languages from around the world (**Figure 2a, Supplementary Table 4**).

In order for a group of phages to be deemed a defined lineage, they need to be a monophyletic group that is distinct from established taxonomy^27^. To establish this, we performed proteomic-based analyses on the 1,237 complete genomes against phages classified by the ICTV. In a genomic protein-sharing network, our genomes formed a cluster separate from RefSeq phages (**Supplementary Figure 2a**). This was confirmed in a phage proteomic tree, were they also formed a separate monophyletic clade distinct from ICTV-recognized sequences (**Supplementary Figure 2b**). Together, these results show that the *Ca. Heliusviriales* are a cohesive lineage that is distinct from the current ICTV-ratified families.

A genomic network of the *Ca. Heliusvirales* showed that most subfamilies are highly similar in protein content, a notion supported by conservation of the marker genes and gene synteny, as well as further shared genes between subfamilies, from pairwise comparison between the proposed families and subfamilies (**Figure 3**). Nevertheless, several subfamilies formed distinct clusters in the genomic network, most evidently the *Ca. Mehrvirinae* (*Ca. Aineviridae)* and *Ca. Nitavirinae* (*Ca. Utuviridae)*, and to a lesser extent the *Ca. Mataharivirinae* (*Ca. Hathorviridae*) (**Figure 2c**). This indicates that they are distinct in protein content, which is reflected by the environments in which they were found and their predicted bacterial hosts (**Figure 4a and b**), as well as their divergent GC content when compared to other members of their family (**Supplementary Figure 3**).

**Figure 3:**
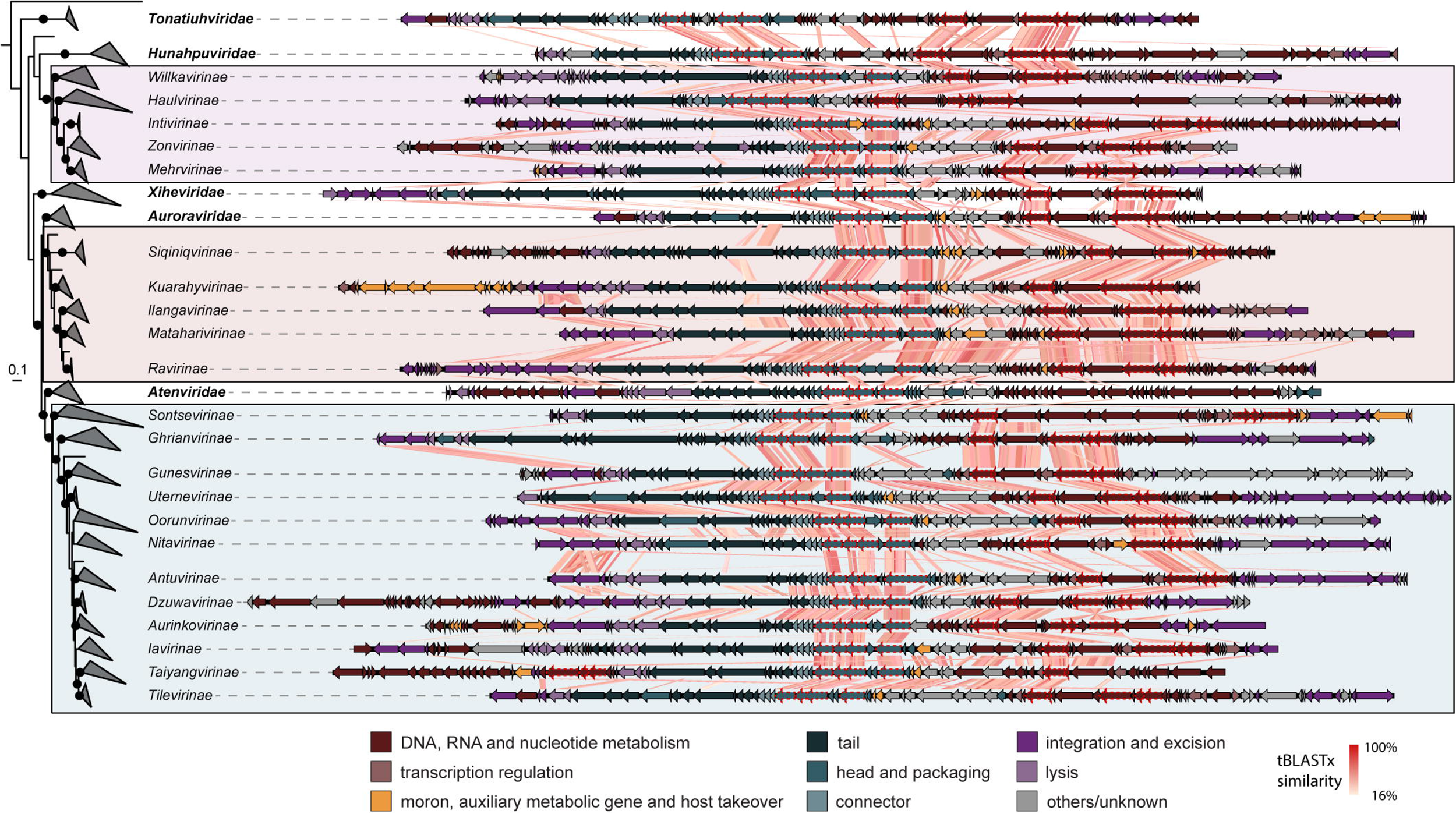
Pairwise whole-genome comparisons of genomes from each of the families and subfamilies, with vertical lines showing tBLASTx similarity. The tree on the left is a collapsed version of the one in Figure 2a, with the outgroup pruned to a single branch (*Rowavirus rowa*). Genes are colored according to their function, and homologs of the nine originally identified marker genes have dashed red outlines. For the names of the genomes depicted, see **Supplementary Table 4**.

**Figure 4:**
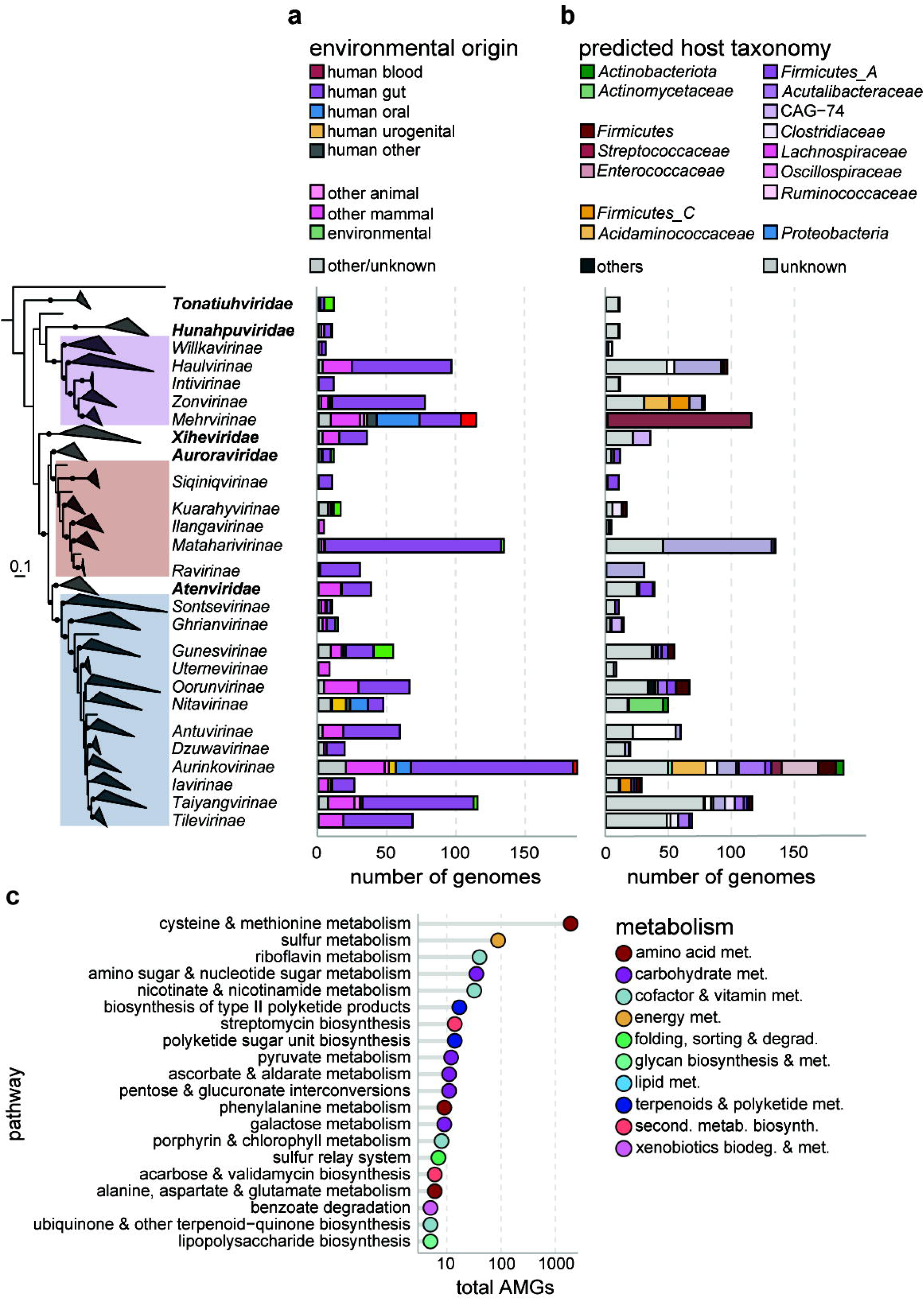
high *Ca. Heliusvirales* diversity in environmental origin and host species. **a:** environmental origins of the *Ca. Heliusvirales* subfamilies. **b:** Iphop-based bacterial host predictions of *Ca. Heliusvirales* lineages. Detailed results can be found in **Supplementary Figure 4** and **Supplementary Table 5. c:** pathway assignment of putative auxiliary metabolic genes (AMGs) and morons encoded by the *Ca. Heliusvirales*. For brevity, the 20 pathways that were most commonly found were selected.

Several subfamilies were distinct in their environmental origins or predicted hosts. The *Ca. Mehrvirinae* were enriched in phages from the human oral cavity (hypergeometric test, *p* = 1.5 × 10^−17^) an environmental preference also reflected in their most commonly predicted hosts: the facultatively anaerobic genus *Streptococcus* (family *Streptococcaceae*). The distinctness of the *Ca. Nitavirinae* similarly reflect their predicted hosts in the *Actinomycetaceae* rather than the *Firmicutes* infected by other *Ca. Heliusvirales*. The *Lachnospiraceae* phages in the *Ca. Mataharivirinae* were enriched in human gut-derived phages (hypergeometric test, *p* = 2.8×10^−21^). Within the *Lachnospiraceae*, the most commonly predicted *Ca. Matahrivirinae* hosts were the genera *Coprococcus*, *Agathobacter, Dorea_A,* and *Ruminococcus_B*, which are all common human gut bacteria (**Supplementary Figure 4**). Finally, the *Ca. Zonviridae*, although less distinct in the genomic network (**Figure 2c**), were peculiar in their predicted hosts. These were in the *Negativicutes*, with their mostly predicted hosts being in the *Acidaminococcaceae* and *Megasphaeraceae* families, which unlike other *Firmicutes* possess diderm cell envelopes^30^. Unlike the recently defined *Crassvirales*, which are all thought to infect hosts within the *Bacteroidetes* phylum^31^, the *Ca. Heliusvirales* thus seem to infect a wide variety of hosts across multiple phyla.

Almost all complete *Ca. Heliusvirales* genomes (92.4%) carried identifiable genes involved with integration and excision, meaning they were temperate phages (**Supplementary Figure 5**). While the remainder of the complete genomes might belong to obligately virulent phages, it might also be that they contain divergent integration genes that evaded annotation or have other chronic lifestyles. The only lineages of which less than 75% or genomes was identifiably temperate contained less than 40 genomes: the *Willkavirinae* (5 genomes), *Xiheviridae* (35 genomes), and *Uternevirinae* (8 genomes), indicating that they are rare. Thus, while *Ca. Heliusvirales* phages are predominantly temperate, eco-evolutionary pressures on phage lifestyle contributed to the divergence of some lineages contained within it.

Putative auxiliary metabolic genes were common among *Ca. Heliusvirales*, being present in 78.1% of genomes (**Figure 4c**), although we found no clear family-or subfamily specific auxiliary metabolic genes. Many were part of cysteine and methionine metabolism, with the most common genes being DNA (cytosine-5)-methyltransferase 1 (K00558, present in 67% of genomes) and S-adenosylmethionine synthetase (K00789, present in 58.9% of genomes). Both of these genes likely aid the phage in defending against bacterial restriction-modification defenses^32,33^. After cysteine and methionine metabolism, sulfur metabolism-associated genes were most commonly found, which could be involved in reprogramming bacterial sulfur metabolism for additional energy production during phage particle formation^34^.

### Helius-phages have been associated with humans since ancient times

Across our datasets, we found *Ca. Heliusvirales* TerL genes on every continent (**Figure 5a**), indicating that they are widespread. Only one country had human gut-associated samples without any detected *Ca. Heliusvirales*: El Salvador (n=1 sample). Indeed, analysis of 7,166 human gut metagenomes of 5,441 individuals from 38 studies^35^ found *Ca. Heliusvirales* TerL genes in 4,491 individuals (80%, **Figure 5b**). Outside the gut, prevalence of *Ca. Heliusvirales* was markedly lower: we detected their presence in 18/54 skin samples, 17/61 vaginal samples, and 115/312 oral samples. While the *Ca. Heliusvirales* thus occur in various body sites, they are especially prevalent in the gut. This is in line with the finding that their most commonly predicted hosts are in the *Firmicutes* (e.g., the *Lachnospiraceae*), which is one of the main bacterial phyla in the human gut.

**Figure 5:**
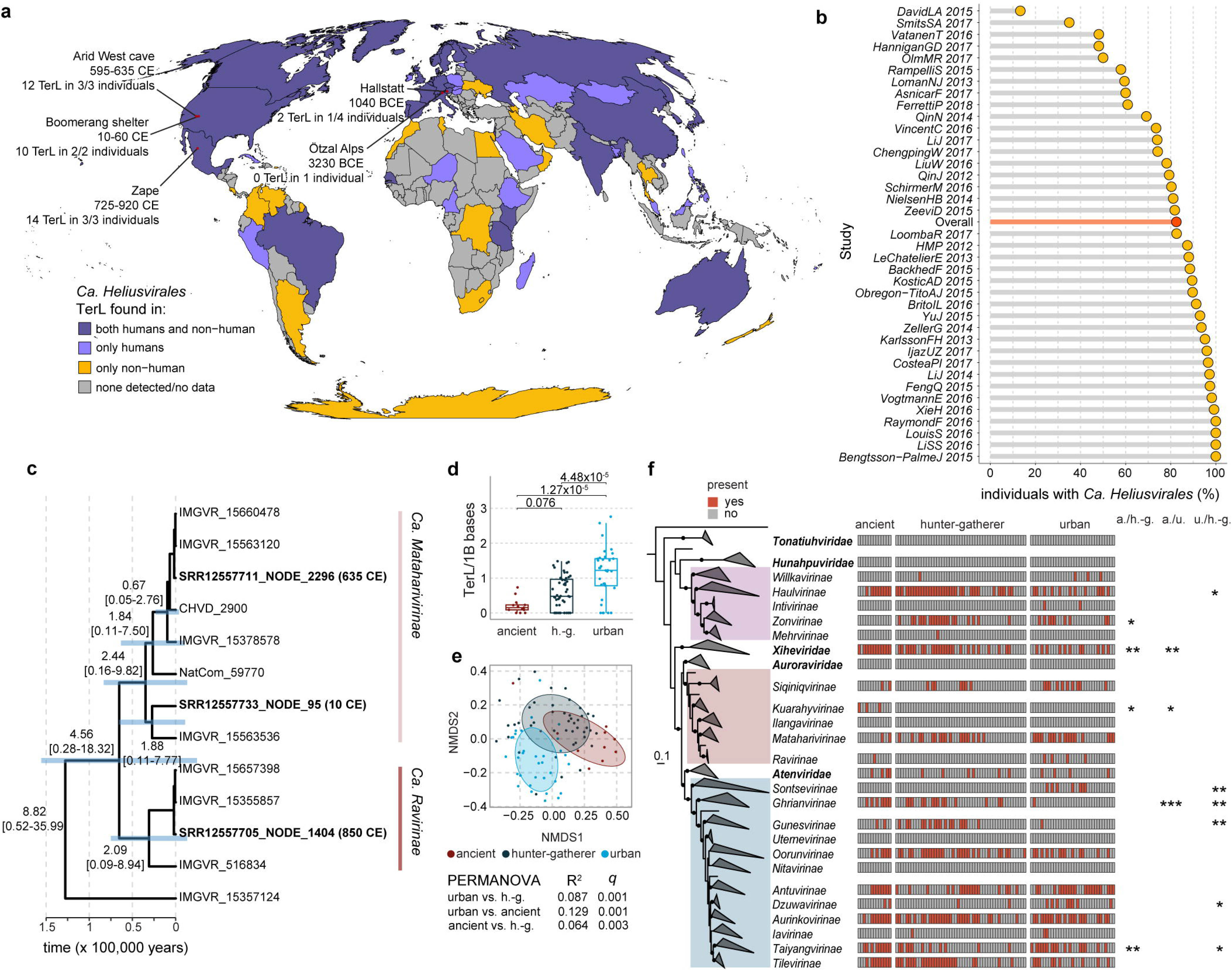
the *Ca. Heliusvirales* are widespread and ancient. **a:** map showing in which countries *Ca. Heliusvirales* TerL-containing sequences were found. Data pertains to the seven bacteriophage databases described in the methods and samples assembled earlier by Pasolli *et al*^35^. Red dots show the locations of ancient gut microbiomes that were analyzed for *Ca. Heliusvirales* TerL presence, with their dates, gene-, and sample-counts. **b:** prevalence of *Ca. Heliusvirales* TerL genes among individuals of 39 gut metagenome studies. The red bar is the prevalence among the totality of all studies. **c:** tip-dated phylogenetic tree of selected *Ca. Hathorviridae* sequences from modern and ancient samples. Bold sequences are from ancient samples. Bar plots denote 95% confidence, values denote median and 95% confidence intervals. A phylogenetic tree of all *Ca. Hathorviridae* sequences is shown in **Supplementary Figure 6**. **d:** boxplots of the number of *Ca. Heliusvirales* TerL genes found per person, per billion sequenced bases, with jittered points per study. Significance is according to nested linear mixed-effect models where the study was used as a random effect. Box plots show the median (middle line), 25th, and 75th percentile (box), with the 25th percentile minus and the 75th percentile plus 1.5 times the interquartile range (whiskers). Comparisons of richness in urbanized vs hunter-gatherer populations within each study can be found in **Supplementary Figure 7a**. **e:** non-metric multidimensional scaling (NMDS) on unweighted UniFrac distances, colored by lifestyle of the human host. **f:** Prevalence of *Ca. Heliusvirales* subfamilies in ancient peoples, hunter-gatherers, and urbanized peoples, with significance according to Fisher exact tests for subfamilies with significant values. All p-values were adjusted for multiple testing with the Benjamini-Hochberg procedure, and are denoted as follows * ≤0.05, ** ≤0.01, *** ≤0.001.

We hypothesized that the high *Ca. Heliusvirales* prevalence reflects its ancient association to the human lineage. To explore this, we re-analyzed whole metagenome shotgun sequencing datasets from paleofeces in Austria^36^ and North America^37^. Among the North-American samples, which were from three different locations across the USA and Mexico, and were dated between 10-920 CE, we found 36 *Ca. Heliusvirales* TerL genes across all eight of the studied samples (**Figure 5a**). In the Austrian samples, we found four TerL sequences in one of the four Austrian samples, which was dated at between 652-544 BCE. We furthermore analyzed gut and small-intestinal samples from an ancient European individual who lived around 3,200 BCE^38^, but found no clear *Ca. Heliusvirales* sequences, although four genes with hmmsearch bit-scores of between 600 and 700 against the TerL were present. These samples may thus contain distant relatives of the *Ca. Heliusvirales*. The greater absence of *Ca. Heliusvirales* among the European samples may reflect higher levels of degradation or fragmentation of these samples.

The presence of *Ca. Heliusvirales* in multiple pre-colonial North Americans implies that they were part of the human gut microbiome before human migration to the Americas (about 15,000 years ago^39^). We confirmed this hypothesis with a time-measured phylogenetic tree of the two main *Ca. Hathorviridae* subfamilies, which contains the strongly human-gut associated *Ca. Mataharivirinae*. A tree built with an optimized relaxed clock of ancient North-American sequences and a selection of modern human gut sequences showed that both the *Ca. Mataharivirinae* and *Ca. Ravirinae* started diversifying about between 210,000 and 250,000 years ago, though with large 95% highest posterior density (HPD) intervals (*Ca. Mataharivirinae*:244,143 years ago (95% HPD 15,000-981,000), *Ca. Ravirinae*: 209,075 years ago (95% HPD 9,200-894,000), **Figure 5c**) This is around the time that *Homo sapiens* first emerged^40^, clearly indicating that these phages have been a part of the human gut ecosystem since our distant past.

Ancient gut microbiomes have similar levels of bacterial diversity to those of modern populations consuming non-urbanized diets^37^, which are more diverse than those consuming urbanized diets^41,42^. By extension, this is likely true for the gut viruses as well. To further explore the relation between ancient, non-urbanized and urbanized *Ca. Heliusvirales*, we compared them to samples derived from urbanized and non-urbanized populations. To minimize inter-study batch effects, we selected two studies that included both hunter-gatherers (from Tanzania or Peru) and urbanized people (from Italy or the USA)^41,42^. Within both studies, we observed a significantly higher *Ca. Heliusvirales* richness among urbanized people (Wilcoxon signed-rank test, Benjamini-Hochberg-adjusted *p*=0.026 and 0.01, **Supplementary Figure 7a**). Combining the values of modern urbanized and hunter-gatherer populations with ancient ones showed significantly higher *Ca. Heliusvirales* richness in urbanized people than both hunter-gatherers (linear mixed-effect model, Benjamini-Hochberg-adjusted *p*=4.5×10^−5^) and ancients (linear mixed-effect model, Benjamini-Hochberg-adjusted *p*=1.2×10^−5^), but not between hunter-gatherers and ancients (linear mixed-effect model, Benjamini-Hochberg-adjusted *p*=0.076, **Figure 5d**). This interestingly is the reverse of overall gut bacterial richness, which decreases with urbanization^43^. Abundance, measured as the fraction of reads that mapped to *Ca. Heliusvirales* TerL genes, was not significantly different between the three populations (**Supplementary Figure 7b**). *Ca. Heliusvirales* thus seem to diversify in tandem with urbanization.

We next analyzed *Ca. Heliusvirales* β-diversity among urbanized, hunter-gatherer, and ancient samples with a phylogenetic tree of TerL genes from these samples and those from complete genomes as depicted in Figure 2a. Non-metric multidimensional scales (NMDS) on unweighted UniFrac distances intriguingly showed that all three populations (urban, hunter-gatherers, ancients) had distinct *Ca. Heliusvirales* populations (permanova *q*=0.001, **Figure 5e**). Among ancient samples the small *Ca. Xiheviridae* family were particularly more prevalent than both modern hunter-gatherers and urban populations (Fisher’s exact test, Benjamini-Hochberg-adjusted *p<*0.05, **Figure 5f**). The *Ca. Ghrianvirinae* (*Ca. Utuviridae*) were meanwhile much more prevalent among both ancient and modern hunter-gatherers than urbanized populations. In general, the number of observed *Heliusvirales* subfamilies per billion sequenced bases was the lowest among ancient samples, with no discernable difference between modern hunter-gatherers and urban populations (**Supplementary Figure 7c**). This indicates that modernity has led to an expansion of *Ca. Heliusvirales* that is not only explained by life style. Our results could reflect that *Ca. Heliusvirales* hosts in the *Lachnospiraceae* thrive in the urbanized gut microbiome.

### Helius-phage richness is associated with disease

Our previous research^8^ focused on gut virome alterations in metabolic syndrome (MetS), and found an association between members of the *Ca. Heliusvirales* and this set of cardiometabolic risk factors. To study a wider range of illnesses, we now identified *Ca. Heliusvirales* sequences in fourteen studies of diverse human-derived samples^35^ by searching for the TerL marker gene. Within-study analyses identified significantly altered *Ca. Heliusvirales* richness in four out of twelve illnesses (Wilcoxon signed-rank test Benjamini-Hochberg adjusted p<0.05): type 1 diabetes (T1D), type 2 diabetes (T2D), inflammatory bowel disease (IBD), and liver cirrhosis (**Figure 6a).** In the former three, richness was elevated when compared to healthy controls, while in liver cirrhosis it was decreased. These disease linkages are interesting in light of the predicted *Lachnospiraceae* hosts of many *Ca. Heliusvirales*, because this bacterial family includes species involved in short-chained fatty acid production (e.g., *Roseburia intestinalis, Anaerobutyricum hallii, Coprococcus eutactus,* and *Blautia wexlerae* **Supplementary Figure 4**)^44^, which are often associated with the healthy gut. Indeed, increased *Lachnospiraceae* abundance in T1D-^45^ and T2D-patients^46^ has been reported, though a study of *in vitro* incubations in a relatively small cohort size (n=26) linked IBD to decreased *Lachnospiraceae* abundance^47^. For the former two diseases, this could explain the greater richness and prevalence in *Ca. Heliusvirales* phages among such patients. The alterations in the *Ca. Heliusvirales* order are notably counter to ASV-level analysis of the entire gut virome, which was found to be unchanged in T1D^48^, T2D^49^, and IBD^1^. This underscores the value of analyzing higher taxonomic ranks in the context of the gut virome.

**Figure 6:**
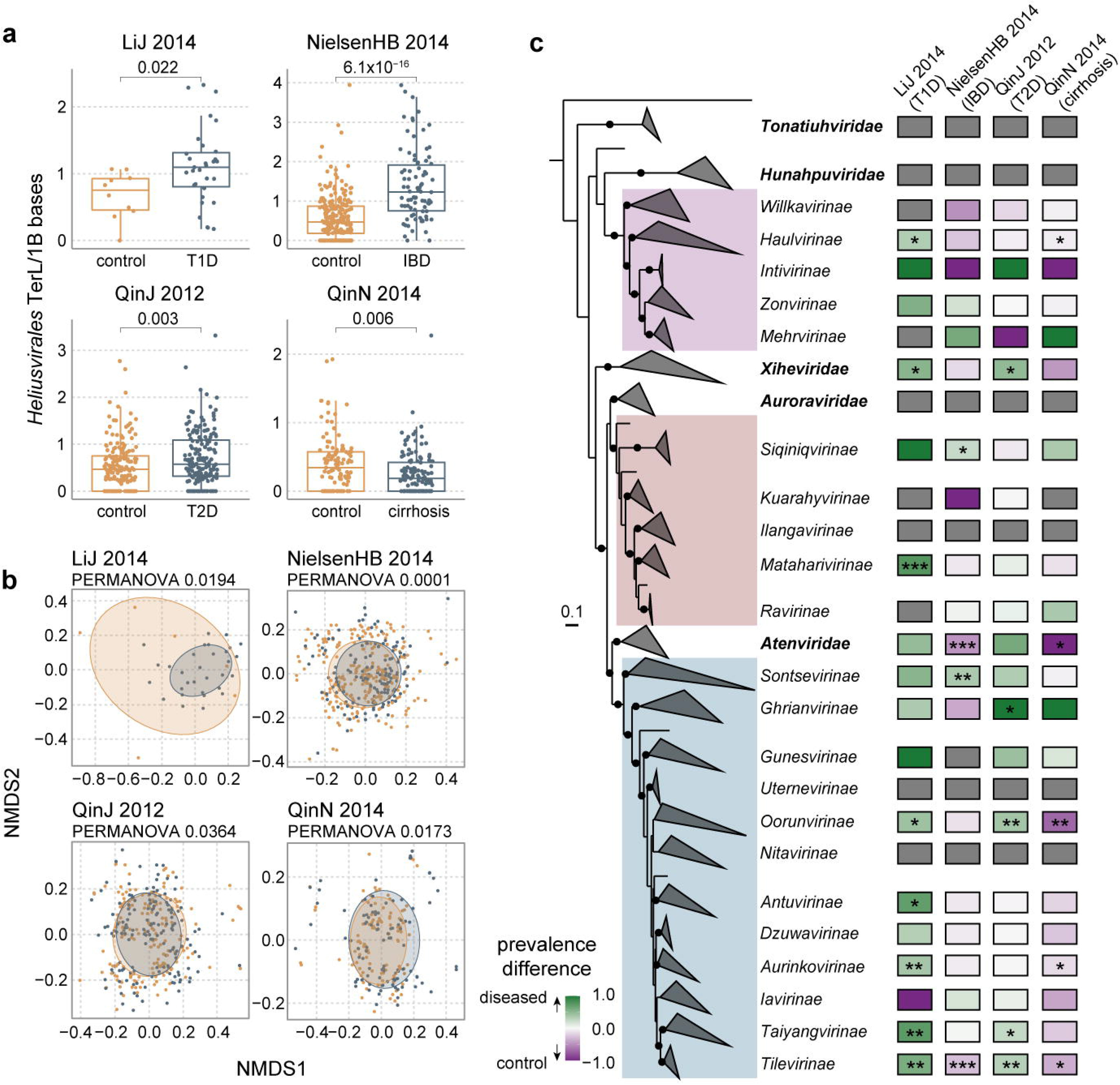
*Ca. Heliusvirales* phages are associated with a number of diseases. **a:** four studies in which a significant difference was found in the number of *Ca. Heliusvirales* TerL genes per person per billion sequenced bases. Significance is according to Wilcoxon signed-rank tests. Box plots show the median (middle line), 25th, and 75th percentile (box), with the 25th percentile minus and the 75th percentile plus 1.5 times the interquartile range (whiskers). **b:** non-metric multidimensional scaling (NMDS) on unweighted UniFrac distances, colored by disease status. **c:** The difference in prevalence between controls and people with disease. All p-values were adjusted for multiple testing with the Benjamini-Hochberg procedure.

Besides differences in *Ca. Heliusvirales* richness, the populations of these phages also differed in each of the four studies (unweighted UniFrac, PERMANOVA Benjamini-Hochberg adjusted *p*<0.05, **Figure 6b**). These population-wide differences between healthy controls and people with T1D, T2D, IBD, and cirrhosis expressed themselves in distinct patterns of prevalence at the subfamily level (**Figure 6c**). Similar to differences between ancient, modern hunter-gatherer, and modern urbanized populations (**Figure 5f**), the *Ca. Xiheviridae* phages were more prevalent in T1D- and T2D-patients than the corresponding healthy individuals (Fisher’s exact test, Benjamini-Hochberg adjusted *p*<0.05), though not in IBD or cirrhosis. The *Ca. Ghrianvirinae*, which was less prevalent in modern urbanized populations than either modern hunter gatherer or ancient ones, was more prevalent in T2D patients than controls, but not in the other diseases. Besides these two, the *Ca. Tilevirinae*, was significantly more present in T1D and T2D than controls, and significantly less so in IBD and cirrhosis. The fact that similar *Ca. Heliusvirales* were depleted in both urbanized versus non-urbanized populations and T1D/T2D patients versus healthy controls perhaps indicates that they are the viral component of VANISH (volatile and/or associated negatively with industrialized societies of humans) gut microbiome taxa^50^.

## Discussion

Elucidating higher-order structures among gut phage lineages is essential for gaining a deeper understanding of the viromes in the human gut and beyond. This process is underway with the re-evaluation of viral taxonomy along genomic lines^27^. Recently, the prevalent *Crassvirales* phage order has undergone the arduous process of classification^31,51^, and here we add the *Ca. Heliusvirales* order as a second genome-based classification of phages that are widespread in the human gut. For this, we used a combination of marker gene phylogenies, genomic analyses, and proteomics-based approaches. As the TerL marker gene is conserved among the *Duplodnaviria*, to which dsDNA bacteriophages belong^52^, we considered the phylogeny built with this gene is the most reliable method for defining *Ca. Heliusvirales* taxonomy. Our usage of the TerL gene to define taxonomy is akin to the *Crassvirales*^12^, but unlike the concatenated phylogeny of marker genes used to define the *Herelleviridae*^13^. The absence of any other universal marker gene beside the TerL gene among the *Ca. Heliusvirales* invalidates a concatenated phylogeny approach. This absence of universal marker genes also indicates high levels of genomic mosaicism in the *Ca. Heliusvirales*, making them interesting targets to study evolutionary dynamics of human gut viruses.

The *Ca. Heliusvirales* order is highly diverse in environmental origin, geographic distribution, and bacterial hosts. While we found *Ca. Heliusvirales* phages in terrestrial, marine, and gut microbiomes of both cattle and wild animals, the majority of genomes derived from human gut samples. While at face-value this may be assigned to anthropocentric sampling bias, this is unlikely because the IMG/VR database included in our analysis contains about six times more aquatic and terrestrial phages than human-derived ones^53^. Furthermore, non-animal-associated *Ca. Heliusvirales* phages are largely constrained to specific families, indicating environmental differentiation.

*Ca. Heliusvirales* phages associated with the human gut were identified in countries from every permanently inhabited continent, and in about 80% of the population. This percentage could well be an underestimation, as shallower sequencing depths may lead to false negatives. In either case, this order is clearly a core component of the human gut virome. This has been the case since the deep past of modern humans, as evidenced by the presence of these phages in palaeofaeces and our accompanying evolutionary studies. We dated the diversification of the *Ca. Mataharivirinae* and *Ca. Ravirinae* subfamilies roughly to the emergence of *Homo sapiens* in Africa between 200,000 and 300,000 years ago^40^. This dating is notably more ancient than the previously studied diversification of the human-associated bacterial species *Methanobrevibacter smithii*^37^ and *M. oralis*^54^ at respectively 85,000 and 126,000 years ago. This is likely due to the differing taxonomic levels at which analyses where done, which we were forced to do due to the relative sparce number of phages from the same species among the ancient samples. Additional ancient samples might quell these issues and allow for an analysis at lower taxonomic levels. Furthermore, due to the high sequence diversity in our timed phylogenetic tree, we obtained rather large 95% confidence intervals, which are also likely to shrink as further ancient microbiome data becomes available. During their rapid evolution, these phages perhaps diversified in response to microbiome alterations influenced by changing human geography and diet, both main drivers of human microbiome composition^55,56^. Earlier studies showed co-linearity of *Crassvirales* between humans and great apes^57^, and estimated a diversification of these phages in the past several centuries^58^. This estimate of much more recent diversification was, however, noted for its uncertainty and focused only on *crAssphage* rather than the subfamily-level analysis presented here.

We also found that higher *Ca. Heliusvirales* richness is associated with increased urbanization. This is remarkable, as microbiome diversity as a whole is lower with increasing urbanization^41,42^. The opposite observations in *Ca. Heliusvirales* could reflect the success of their bacterial hosts to replicate in urbanized microbiomes: indeed, Hadza hunter-gatherers from Tanzania and Matses hunter-gatherers from Peru have lower levels of *Lachnospiraceae* bacteria such as *Blautia* and *Ruminococcus* than people in urbanized environments^42,59^.

While the *Ca. Heliusvirales* as a whole are near universally present in the human gut, no single viral genome in this putative order is. The linkages to lifestyle and disease demonstrated here are thus only evident when studying higher viral taxonomic levels. In line with the recent reassessment of viral taxonomy along genomic lines^12^, this study thus shows how future definition of additional viral orders and classes can provide key insights into the ecology and disease-associations of the gut virome.

## Methods

### Data

All bioinformatic tools were used with default settings, unless explicitly stated otherwise. Statistical analyses were done in R v4.2.1. All p-values were adjusted for multiple testing with the Benjamini-Hochberg method.

The primary dataset of *Heliusviridae* marker genes were derived from 298 contigs that were used in our earlier study^8^, which studied the gut virome in the general population-based multi-ethnic Healthy Life in an Urban Setting (HELIUS) cohort^60^. Contig datasets that were used to identify *Heliusviridae* consisted of 63,481 genomes from this previous study. This included all viral contigs before removing redundant ones. Further data sources consisted of four recently constructed databases of gut bacteriophages: the gut phage database (GPD, 142,809 genomes)^18^, cenote-taker 2 human virome database (CHVD, 93,860 genomes)^20^, gut virome database (GVD, 33,242 genomes)^19^, and metagenomic gut virus catalogue (MGV, 189,680 genomes)^61^. Further contigs were derived from fourth version of the integrated microbial genomes viral dataset (IMG/VR, 15,722,824 genomes)^21^ and bacterial viruses from the NCBI viral reference sequence dataset (RefSeq, 4,220 genomes) release 216. For studies of urbanization- and disease-links, metagenome assemblies were obtained from Pasolli *et al*^35^. of 7,166 human gut metagenomes of 5,441 individuals from 38 studies.

### Identification of putative Heliusviridae sequences

Since previous findings showed *Heliusviridae* genomes to be around 50,000-100,000 bp long, we first selected contigs of at least 25,000 bp without any ambiguous bases from all datasets. This resulted in an effective search space of 1,314,988 contigs. Open reading frames (ORFs) were then predicted using prodigal v2.6.3^62^ (option --meta).

For each of the nine *Heliusviridae* marker genes, a profile hidden Markov model (HMM) was constructed from protein sequences derived from our previous study using hmmbuild v3.3^63^. The resulting profile HMMs were used to search against predicted ORFs using hmmsearch v3.3^63^. For phylogenetic studies, we selected contigs with an hmmsearch hit (bit-score ≥600) against the *Heliusviridae* terminase large subunit (TerL). Duplicates were removed with dedupe from BBTools v38.84 (option minidentity=100), resulting in a non-redundant dataset of 6,359 contigs. These were then analyzed for completeness with checkv v1.0.1^22^ (using database version 1.4), and we selected contigs that had 100% completeness, no warnings, and were not recognized as prophages. These genomes were fully annotated with pharokka v1.2.1^24^ (options -m, -g prodigal), and their ORFs were grouped in protein clusters using mmseqs2 v14.7e284^64^. GC contents were derived from pharokka output, and temperate phages were defined as those with at least one ORF with a predicted function in the “integration and excision” category. Potential auxiliary metabolic genes encoded by the genomes were identified with VIBRANT v1.2.1^65^.

### Viral sequence clustering and proteomic analysis

Separation between our phages and those in the NCBI refseq database was analyzed with both ViPTreeGen v1.1.3^66^ and vContact2 v0.11.3^67^. The vContact2 network was visualized with Cytoscape v3.9.1^68^, while the ViPTreeGen tree was visualized with the interactive tree of life (iTOL) webtool^69^. To analyze the grouping of genomes among the high quality contigs, we used them to constructed a second vContact2 network.

### Phylogenetic tree construction

Before phylogenetic tree construction, incomplete protein sequences were removed by selecting only sequences that were longer than the 25^th^ percentile minus 1.5 times the inter-quantile ranges of all lengths. As outgroup, the homologous TerL sequence from *Staphylococcus* phage LH1 (NC055041) was added. Marker genes were aligned using MAFFT v4.753 (options --maxiterate, --localpair), after which positions with more than 90% gaps were trimmed with trimal v1.4.rev15^70^ (option -gt 0.1). Finally, a tree was built using IQTree v2.2.0.3^71^, using model finder^72^ and 1000 iterations of the ultrafast bootstrap approximation^26^ and SH-like approximate likelihood ratio test. IQTree performed ten separate tree iterations and selected the one with the best log likelihood score (options -B 1000 -alrt 1000 --runs 10). The tree was visualized with the interactive tree of life (iTOL) webtool^69^.

### Heliusvirales hosts

We predicted the hosts infected by the *Heliusvirales* with iPHoP v1.3.2^73^, where host predictions with a score of at least 90 were considered valid. This resulted in genus-level host predictions for 761 *Ca. Heliusvirales* phages.

### Analysis of ancient samples

For analysis of ancient metagenomes, reads were downloaded from two separate studies^36,37^. These were trimmed with fastp v.0.23.2^74^ (options --detect_adapter_for_pe) and error-corrected with tadpole from the bbmap package v38.90 (https://jgi.doe.gov/data-and-tools/bbtools, options mode=correct, ecc=t, prefilter=2). Error corrected reads were assembled with metaSPAdes v3.15.5^75^. The ancient origins of assembled contigs were determined by mapping reads to contigs of ≥1,000 bp with bowtie2 v2.4.2^76^, and determining C-to-T degradation at the 5’ end of sequencing reads with PyDamage v0.70^77^. Contigs with a predicted accuracy of over 0.5 were considered of ancient origins.

### Heliusvirales prevalence and relation to urbanization

To determine *Heliusvirales* prevalence, we firstly gathered geographic and environmental metadata from all genomes used to build the main phylogenetic tree. Second, contigs obtained from Pasolli et al^35^ were analyzed for the presence of *Heliusvirales* TerL genes. To do this, ORFs were predicted with Prodigal v2.6.3^62^ (option –meta), and used for an hmmsearch with the profile hmm of the TerL constructed from sequences identified in our previous study. Hits with bit-scores of over 600 were selected. The same approach was used to identify *Heliusvirales* TerL genes among ancient populations. Enrichment of phages from certain environments among selected subfamilies was calculated with the phyper function from base R.

To analyze the relation of *Heliusvirales* richness to urbanization, richness was calculated by dividing the number of identified TerL genes by the number of reads in billions. Significance testing of richness differences after combining studies was done with linear mixed effects models where the study was entered as a random effect, using the lme function from the nlme R package v3.1-162 and Anova function from the rstatix R package v0.7.2. The same approach was used to calculate significant richness differences among pooled disease-related cohorts.

Assignment of TerL sequences from ancient- and disease-related samples was done by constructing a phylogenetic tree with TerL genes from all high-quality *Heliusvirales* genomes that had been used in the main phylogeny. For the ancient, hunter-gatherers, and urban populations, a single such tree was built, while for the disease-related datasets one tree was constructed per project. Trees were constructed with IQTree v2.2.0.3 as described above, and pairwise cophenetic distances were calculated from them with the ape R package v5.7-1. Each sequence was then assigned to the same taxonomy as its closest neighbor of which the taxonomy was known.

For the ancient, hunter-gatherers, and urban populations, taxonomic assignment at the family level was used to calculate UniFrac distances, from which a non-metric multidimensional scaling (NMDS) was constructed. Permutated ANOVA significance levels were calculated with the adonis2 function in the vegan R package v2.6-4. To determine significant differences in subfamily prevalence between populations with different lifestyles, Fisher exact tests were calculated using the fisher.test function from base R.

### Timed tree construction

To reconstruct an evolutionary timed tree of the *Ca. Hathorviridae*, we first constructed a regular tree on members of this family. For this, we selected the TerL genes of all *Ca. Hathorviridae* from both the complete genomes and the ancient samples. This was used to build a phylogenetic tree with IQTree as described above. From this tree, we selected thirteen complete genomes and all three ancient genome fragments that combined reflected the architecture of the *Ca. Hathorviridae*. All ORFs from these genomes were then clustered with mmseqs2 v14.7e284^64^, after which the six protein clusters that were universally present were selected. This number was relatively low because the ancient sequences were genome fragments of 15,076, 19,664, and 67,881 bp long. The six universal PCs were a TerL, head maturation protease, major head protein, portal protein, a hypothetical protein, and terminase small subunit. For each protein, an alignment was made with MAFFT v4.753 (options --maxiterate, --localpair)^78^, from which a timed tree was inferred using BEAST v2.7.3^79^. BEAST was run with the following models: Birth death model, Coalescent constant population, Coalescent exponential population, Coalescent bayesian skyline, Coalescent extended bayesian skyline, each with both strict and optimized relaxed clock models. Analyses were run for 30 million iterations, at which point all models had converged and estimated sample sized were above 200. The coalescent constant population model with a relaxed clock was the best-fitting model. A full accounting of tree construction with the various evolutionary models is in **Supplementary Table 6**. BEAST was run with OBAMA v1.1.1^80^ for amino-acid model averaging.

## Acknowledgements

PAJ was supported by a Talent Development grant of the Amsterdam Gastroenterology, Endocrinology, and Metabolism research school at the Amsterdam UMC. MN was supported by a personal ZONMW-VICI grant 2020 (09150182010020). BED was supported by the European Research Council (ERC) Consolidator grant 865694: DiversiPHI, the Deutsche Forschungsgemeinschaft (DFG, German Research Foundation) under Germany’s Excellence Strategy – EXC 2051 – Project-ID 390713860, the Alexander von Humboldt Foundation in the context of an Alexander von Humboldt-Professorship founded by German Federal Ministry of Education and Research, and the European Union’s Horizon 2020 research and innovation program, under the Marie Skłodowska-Curie Actions Innovative Training Networks grant agreement no. 955974 (VIROINF). HH was supported by an Aspasia premium (015.017.050) and Dutch Diabetes Research Foundation (2019.82.004). The HELIUS study is conducted by the Amsterdam University Medical Centers, location AMC and the Public Health Service of Amsterdam. Both organizations provided core support for HELIUS. The HELIUS study is also funded by the Dutch Heart Foundation, the Netherlands Organization for Health Research and Development (ZonMw), the European Union (FP-7), and the European Fund for the Integration of non-EU immigrants (EIF). We are most grateful to the participants of the HELIUS study and the management team, research nurses, interviewers, research assistants and other staff who have taken part in gathering the data of this study.

## Author contributions

P.A.d.J. conducted data analysis and wrote the manuscript; B.J.v.d.B., A.H.Z., and M.N. assisted with experimental design and data interpretation; P.A.d.J., B.E.D., and H.H. designed the study. All authors read and provided input on the manuscript.

---

**Supplementary Figure 1:**
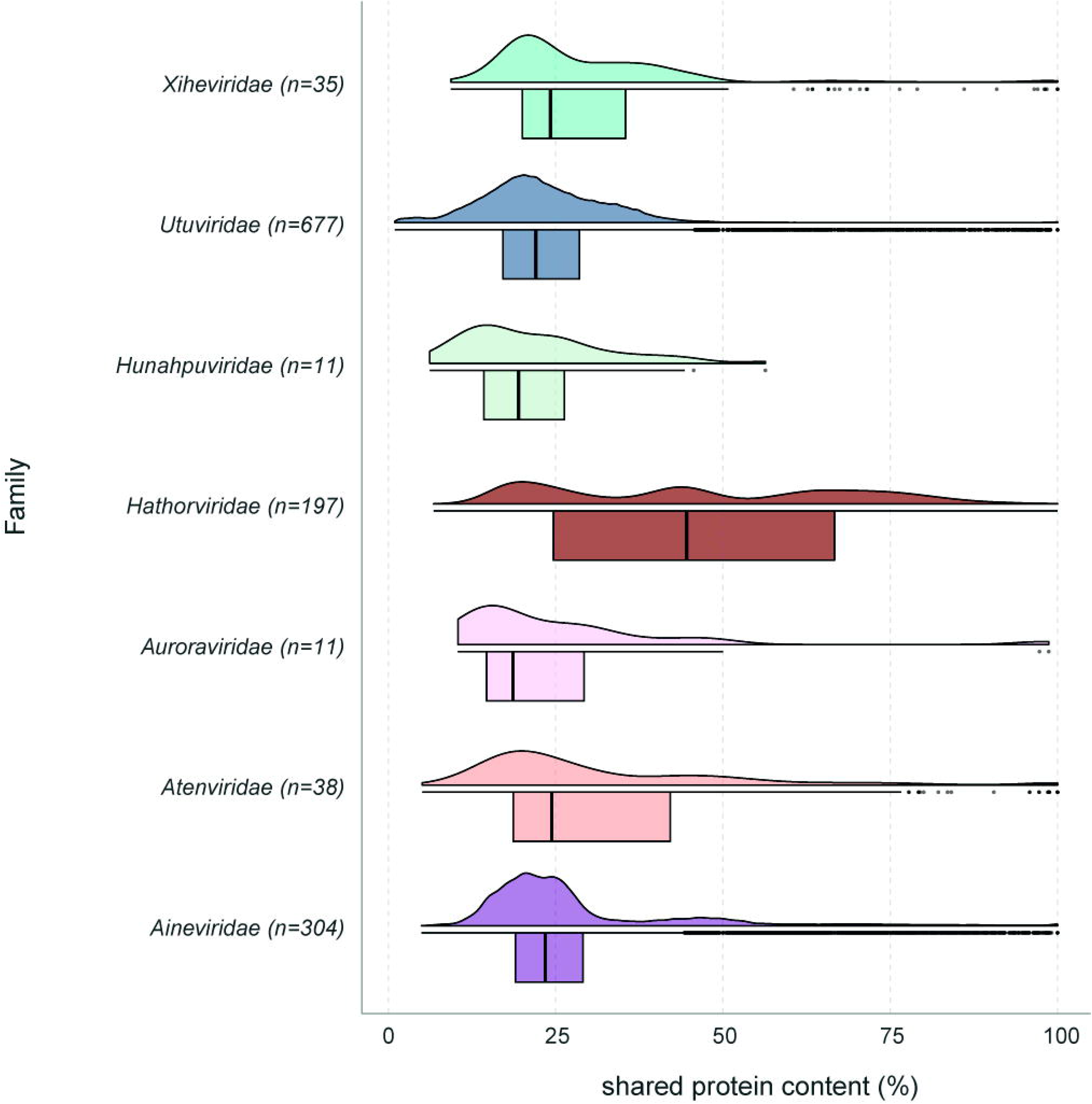
shared gene content for all families.

**Supplementary Figure 2:**
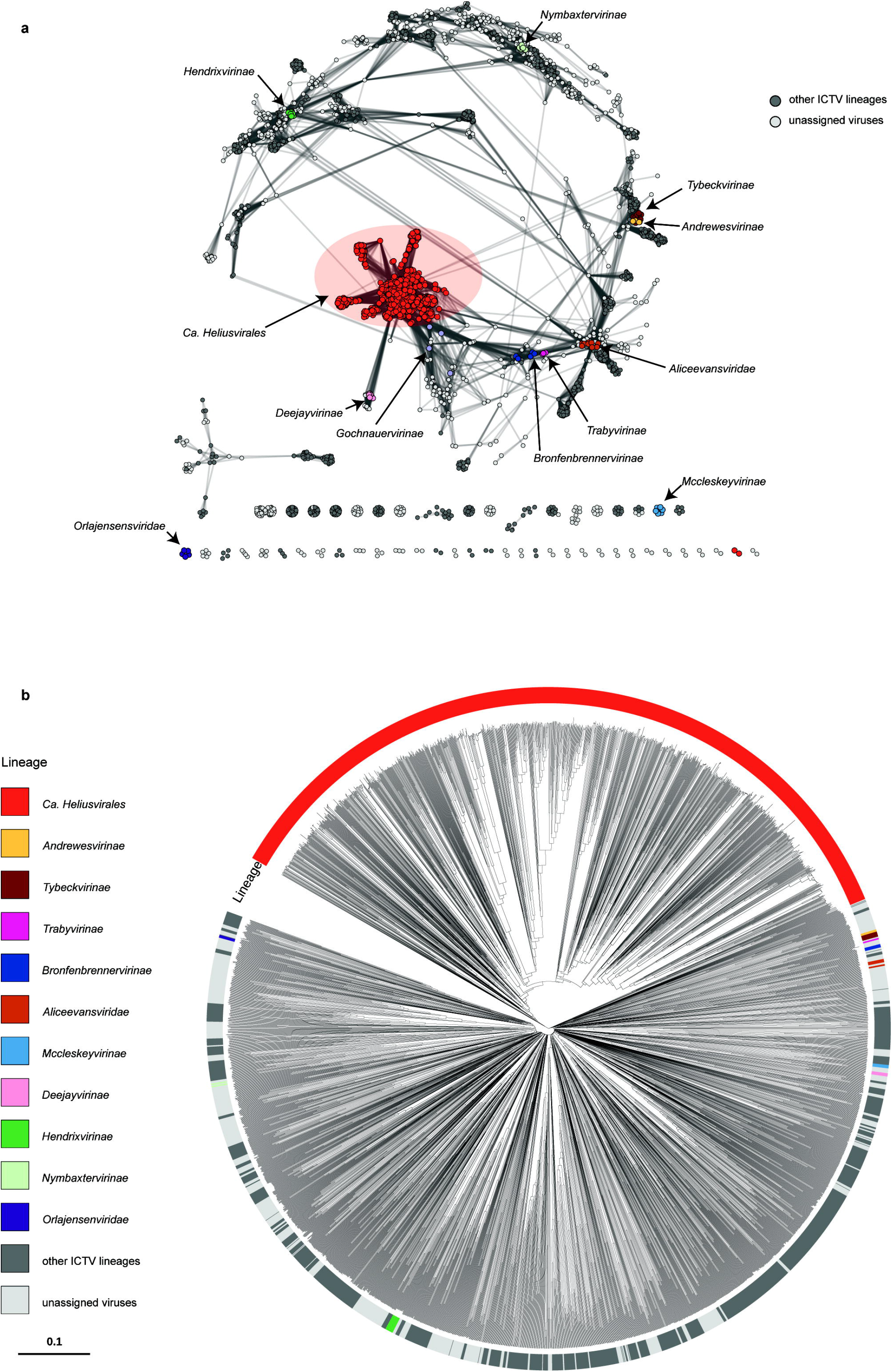
**a**: vContact2 genome network of the *Ca. Heliusvirales* and all genomes in the ICTV database. ICTV lineages that are in Figure 1d are colored **b:** VipTree proteome tree of the same sequences in **a**. Unassigned viruses have no classification above the genus level.

**Supplementary Figure 3:**
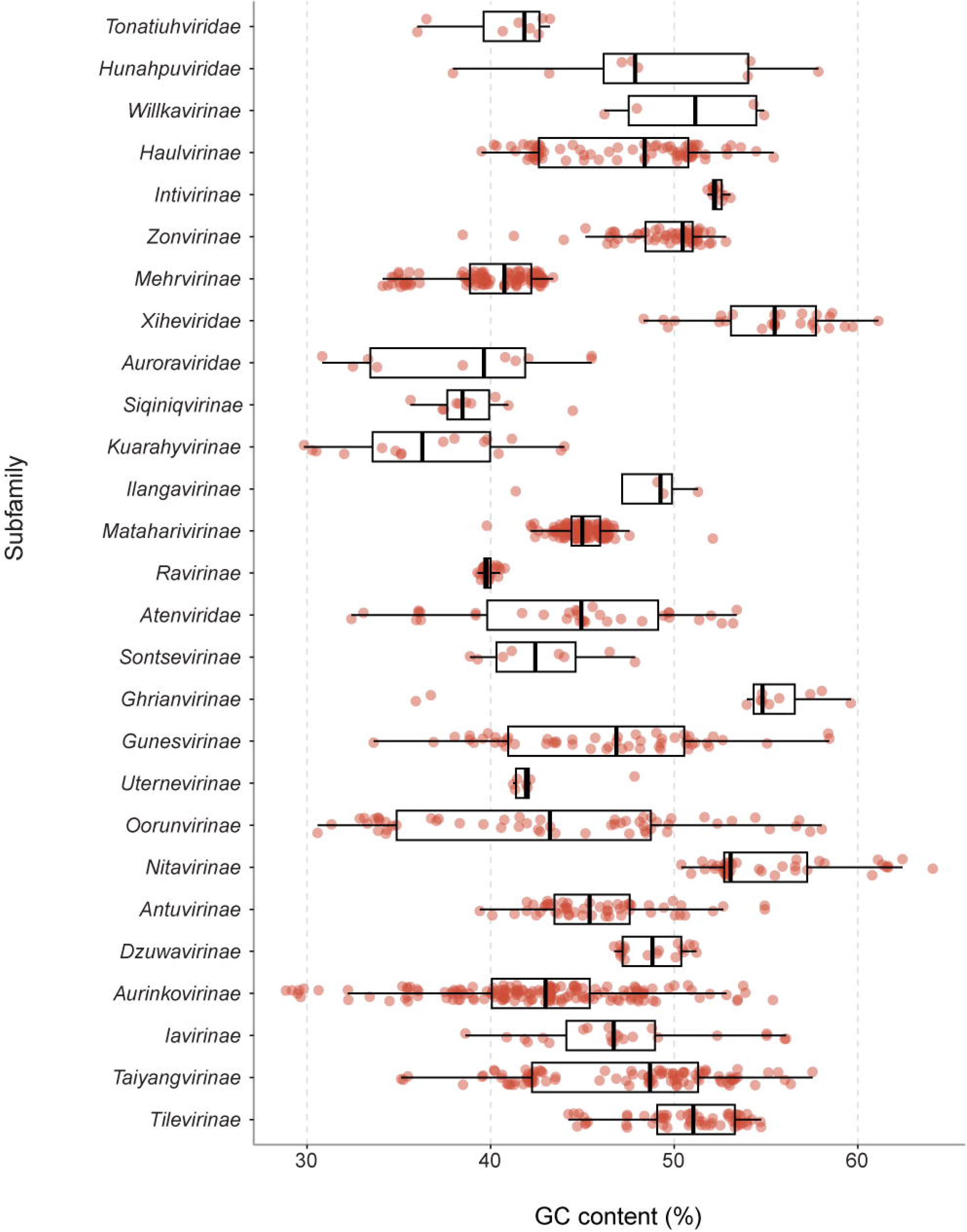
GC-content of the *Ca. Heliusvirales* by lineage.

**Supplementary Figure 4:**
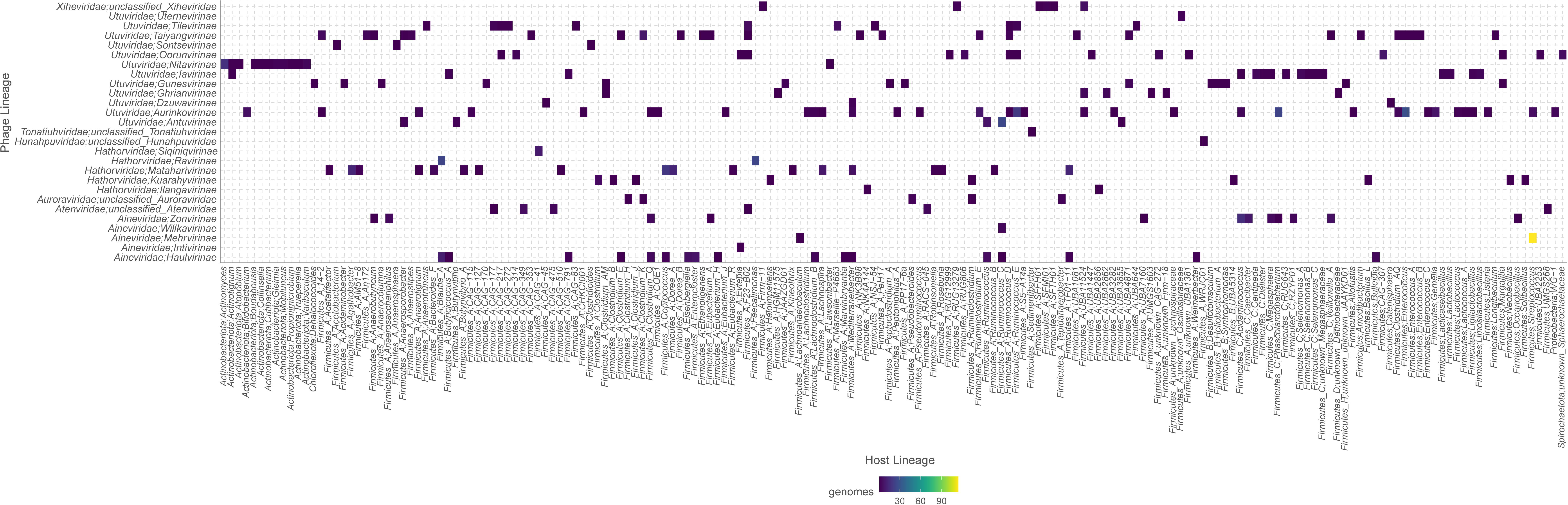
A heatmap of genus-level predictions at the *Ca. Heliusvirales* subfamily-level.

**Supplementary Figure 5:**
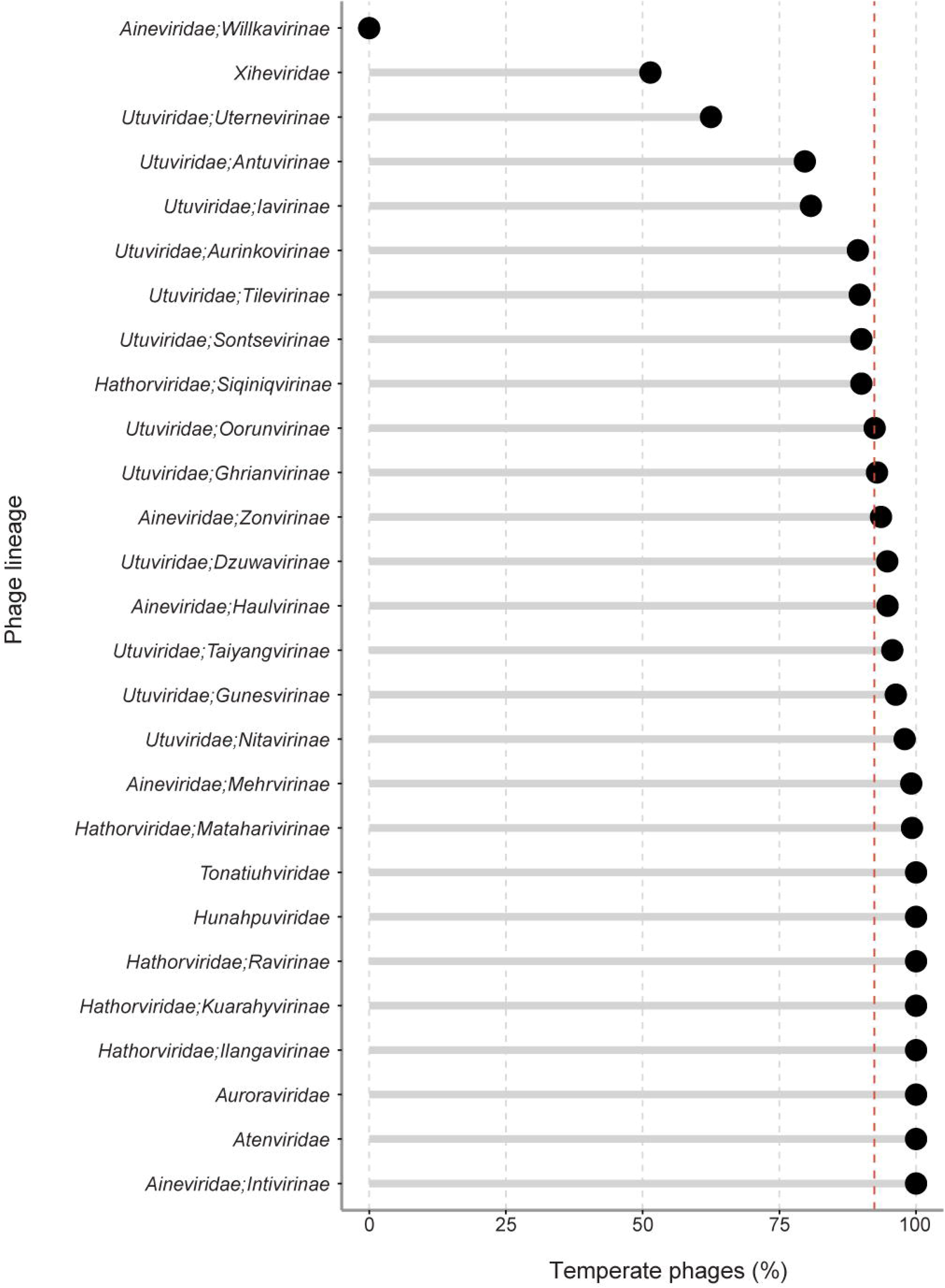
the percentage of temperate phages across the subfamilies, as determined by the presence of integration and excision genes in pharokka annotations. The dashed red line is the average for all *Ca. Heliusvirales*

**Supplementary Figure 6:**
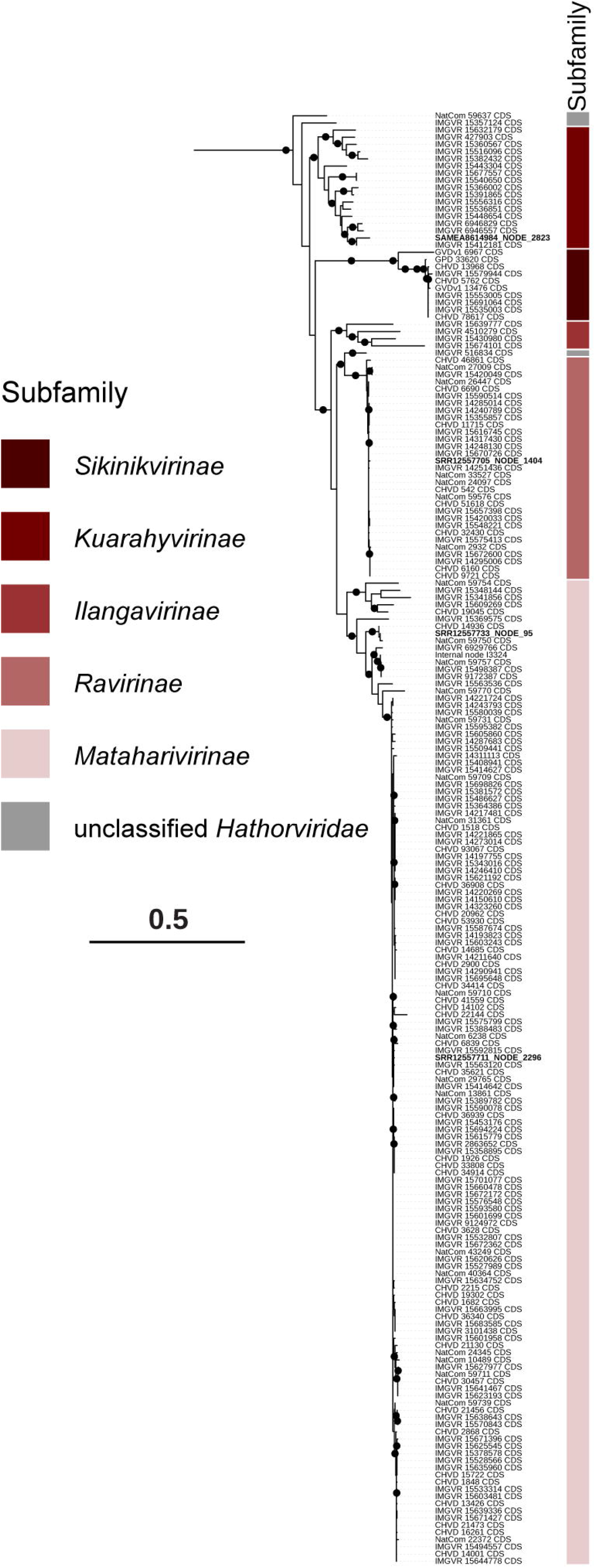
An approximate maximum-likelihood tree of *Ca. Hathorviridae* TerL genes, with color coding showing subfamily. Dots show branches with SH-like approximate likelihood ratio test ≥80 and ultrafast bootstrap approximation ≥95.

**Supplementary Figure 7:**
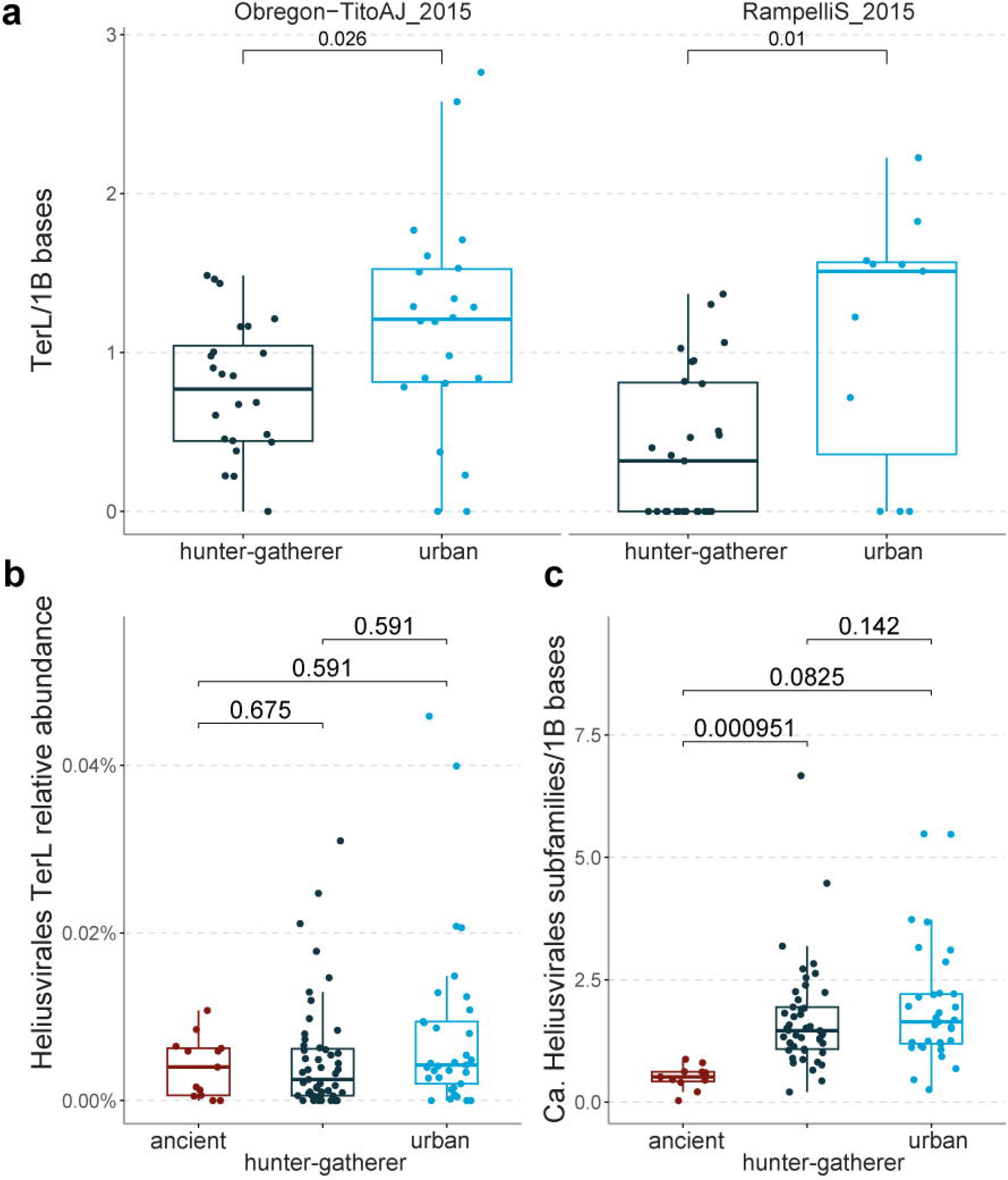
**a**: comparisons of *Ca. Heliusvirales* richness between urban and hunter-gatherer populations in two studies. Benjamini-Hochberg-adjusted p-values are according to Wilcoxon signed-rank tests. **b:** comparison of *Ca. Heliusvirales* TerL abundance between lifestyles. **c:** taxonomic *Ca. Heliusvirales* richness comparison between ancient, hunter-gatherer, and urban populations. Benjamini-Hochberg-adjusted p-values in b and c are according to linear mixed-effect models with the study as a random effect.

